# Identification of the differences in molecular networks between idiopathic pulmonary fibrosis and lung squamous cell carcinoma using machine learning

**DOI:** 10.1101/2024.11.25.625120

**Authors:** Yosui Nojima, Kenji Mizuguchi

**Affiliations:** Artificial Intelligence Center for Health and Biomedical Research (ArCHER), National Institutes of Biomedical Innovation, Health and Nutrition, 17-3 Senrioka-Shinmachi, Settu, Osaka 566-0002, Japan; Center for Mathematical Modeling and Data Science, Osaka University, 1-3 Machikaneyama, Toyonaka, Osaka 560-8531, Japan; Institute for Protein Research, Osaka University, 3-2 Yamadaoka, Suita, Osaka 565-0871, Japan

## Abstract

Idiopathic pulmonary fibrosis (IPF), a form of idiopathic interstitial pneumonia, is an independent risk factor for lung cancer. The prognosis of IPF patients with lung cancer is poorer than that of IPF patients without lung cancer, and preventive measures for lung cancer remain elusive in patients with IPF. To mitigate lung cancer onset in patients with IPF, understanding the distinct mechanisms that induce both diseases is crucial. We developed highly accurate machine learning (ML) models to classify patients with IPF and lung cancer using public RNA sequencing data. To construct the ML models, the random restart technique was applied to five algorithms, namely, *k*-nearest neighbors, support vector machines (SVM) with radial basis function kernel, SVM with linear kernel, eXtreme gradient boosting, and random forest. To identify differentially expressed genes between IPF and lung cancer, feature importance was calculated in the classification models. Furthermore, we detected somatic mutations impacting gene expression using lung cancer data. The ML models identified *VCX2*, *TMPRSS11B*, *PRUNE2*, *PRG4*, *PZP*, *SCARA5*, *DES*, *HPSE2*, *HOXD11*, *S100A7A*, and *PLA2G2A11* as differentially expressed genes. Somatic mutations were detected in four transcription factors, *BHLHE40*, *MYC*, *STAT1*, and *E2F4*, that regulate the expression of the abovementioned 11 genes. Furthermore, a molecular network was discovered, comprising the four transcription factors and 11 downstream genes. The newly identified molecular network enhances our understanding of the distinct mechanisms underlying IPF and lung cancer onset, providing new insights into the prevention of lung cancer complications in patients with IPF.

## Introduction

Idiopathic pulmonary fibrosis (IPF), a common and prognostically unfavorable type of idiopathic interstitial pneumonia (IIP), is characterized by inflammation and fibrotic changes in the lung parenchyma. It features a usual interstitial pneumonia pattern marked by dense fibrotic lesions causing heterogeneous alterations in lung alveolar structures. The pathogenesis of IPF involves lung epithelium or basement membrane injury due to external or internal stimuli on a genetic predisposition background, leading to the proliferation of fibroblasts and myofibroblasts and excessive production of extracellular matrix, thereby disrupting normal lung architecture. This fibrosis progression results in pulmonary sclerosis and respiratory dysfunction (Raghu et al., 2011). IPF, constituting 55%–60% of all IIP cases, has no curative treatment, and the two recommended antifibrotic drugs are only known to slow disease progression (Raghu et al., 2015). IPF frequently coexists with lung cancer (Ozawa et al., 2009). The incidence of lung cancer is 7–14 times higher in IPF cases than in non-IPF cases (Matsushita et al., 1995; Turner-Warwick et al., 1980), and 10%–20% of IPF cases concurrently involve lung cancer (Ozawa et al., 2009). Both conditions are more prevalent in older adults, males, and smokers, sharing risk factors such as smoking and exposure to environmental or occupational hazards (Zaman and Lee, 2018). Additionally, IPF is an independent risk factor for lung cancer.

IPF patients with lung cancer have a significantly worse prognosis than those without lung cancer and lung cancer patients without IPF (Tomassetti et al., 2015; Yoon et al., 2018). Squamous cell carcinoma (SCC) is the most common lung cancer type in patients with IPF, while adenocarcinoma (ADC) is the most common non-small cell lung cancer subtype in the general population (Ballester et al., 2019). Surgical intervention is a treatment option for patients with IPF and lung cancer, but the postoperative incidence of acute exacerbation (AE) of IPF is as high as 10.1% (Sato et al., 2014). Postoperative AE is associated with a mortality of 35.6%–43.9%, indicating a poor prognosis (Sato et al., 2015, 2014). Prevention of lung cancer occurrence is crucial to avoiding AE, but no preventive measures have been established for lung cancer in patients with IPF. Understanding why IPF is an independent risk factor for lung cancer at the molecular level is necessary to prevent lung cancer onset. In contrast to numerous studies on the common molecular mechanisms of IPF and lung cancer onset (Antoniou et al., 2015; Ballester et al., 2019; Leng et al., 2020), research aiming to understand the differences in the onset mechanisms of both diseases is lagging behind.

Surgical intervention in patients with IPF and lung cancer may lead to the risk of AE, making the collection of clinical samples ethically challenging and hindering studies comparing fibrotic and cancerous lungs in these patients. However, Hata *et al*. showed that somatic mutation profiles in SCC are highly similar between patients with or without IPF (Hata et al., 2021), suggesting that genetic information from the cancerous part of IPF patients with lung cancer could be substituted with that from lung cancer patients without IPF. As data on IPF without lung cancer and lung cancer without IPF are relatively abundant in public databases, it is feasible to analyze these datasets to understand the differences between the onset mechanisms of both diseases.

Classical statistics and machine learning (ML) differ in computational tractability with increasing variables per subject. Classical statistical modeling, designed for data with a few dozen input variables and small-to-moderate sample sizes, focuses on unobserved system aspects. Conversely, ML focuses on prediction using general-purpose learning algorithms to search for patterns in complex and unwieldy data (Bzdok, 2017; Bzdok et al., 2018, 2017); these methods can be effective even without a carefully controlled experimental design and amidst complicated nonlinear interactions (Bzdok et al., 2018).

This study uses public RNA sequencing (RNA-Seq) data from the lungs of patients with IPF and SCC, and employs a random restart technique to construct high-accuracy ML models for classifying IPF and SCC. These models help understand the differences in the onset mechanisms of IPF and SCC. The findings of this study highlight the adaptability of omics data to ML, contributing to a better understanding of lung cancer onset and prevention mechanisms in patients with IPF.

## Methods

### Public RNA-Seq analysis and batch effect correction (BEC)

As shown in Table 1, public RNA-seq datasets were obtained from the NCBI Sequence Read Archive (https://trace.ncbi.nlm.nih.gov/Traces/sra/). The quality of the data in the fastq files was confirmed using FastQC (http://www.bioinformatics.babraham.ac.uk/projects/fastqc/). Trimmomatic version 0.36 (http://www.usadellab.org/cms/?page=trimmomatic) (Bolger et al., 2014) was employed to trim the reads using the Illumina TruSeq adapter removal process (2:30:10) and following parameters: LEADING:20, TRAILING:20, SLIDINGWINDOW:4:20, and MINLEN:25. The trimmed reads were then mapped to the reference human genome (version GRCh38) available on Ensembl using HISAT2 version 2.1.0 (http://daehwankimlab.github.io/hisat2/) with default parameters (Kim et al., 2019). The resulting BAM files were input into featureCounts version 2.0.1 (http://subread.sourceforge.net/) (Liao et al., 2014) to generate read count data. Genes listed in Ensembl BioMart (https://www.ensembl.org/info/data/biomart/index.html) as protein-coding genes, lncRNA, or miRNA were selected for further analysis using the biomaRt package (version 2.54.1) in R version 4.2.3 (https://www.r-project.org/). The count data were then converted into transcripts per million (TPM), and the TPM values were subsequently transformed into log_2_ (TPM + 1).

**Table 1.**
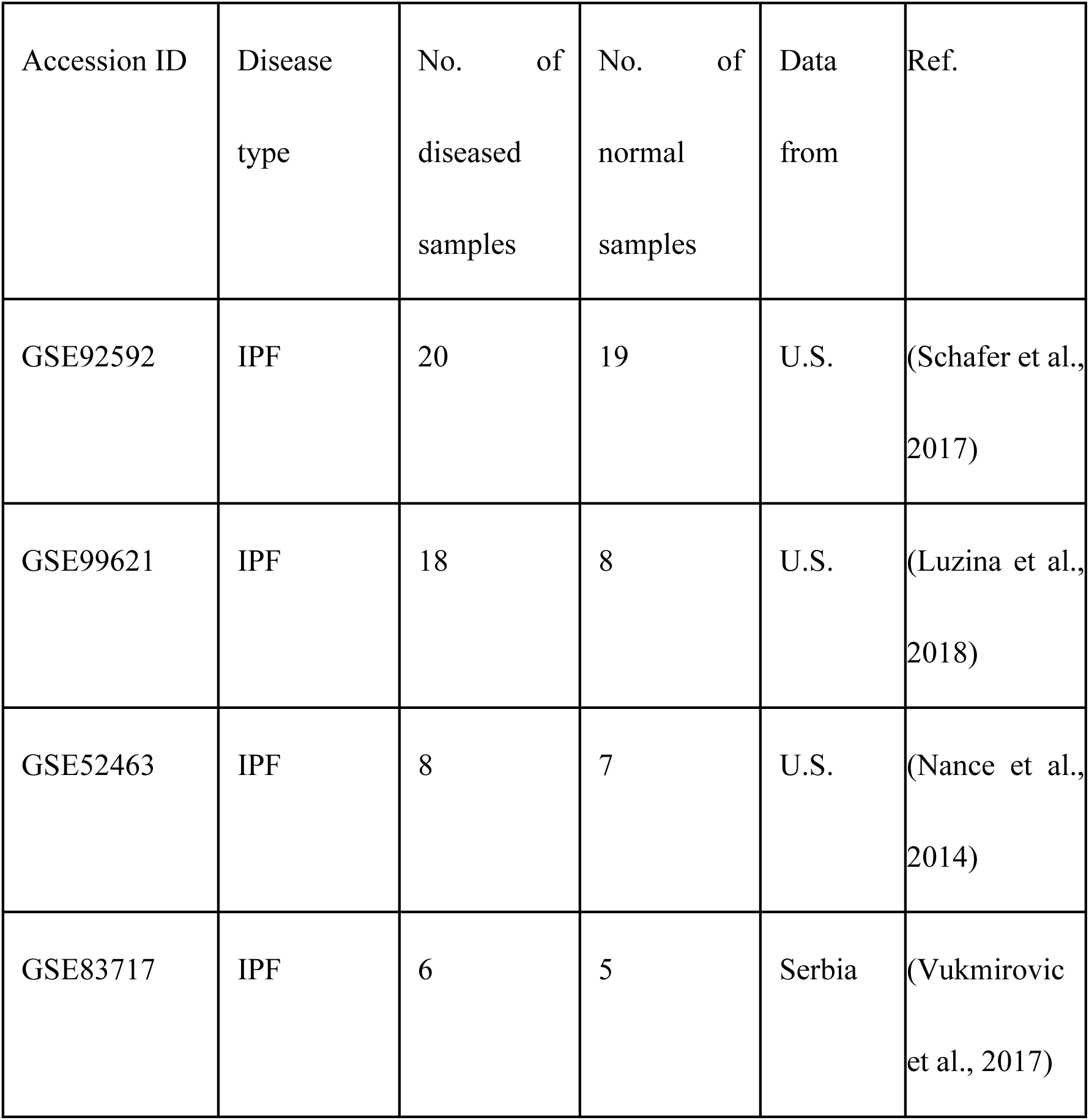
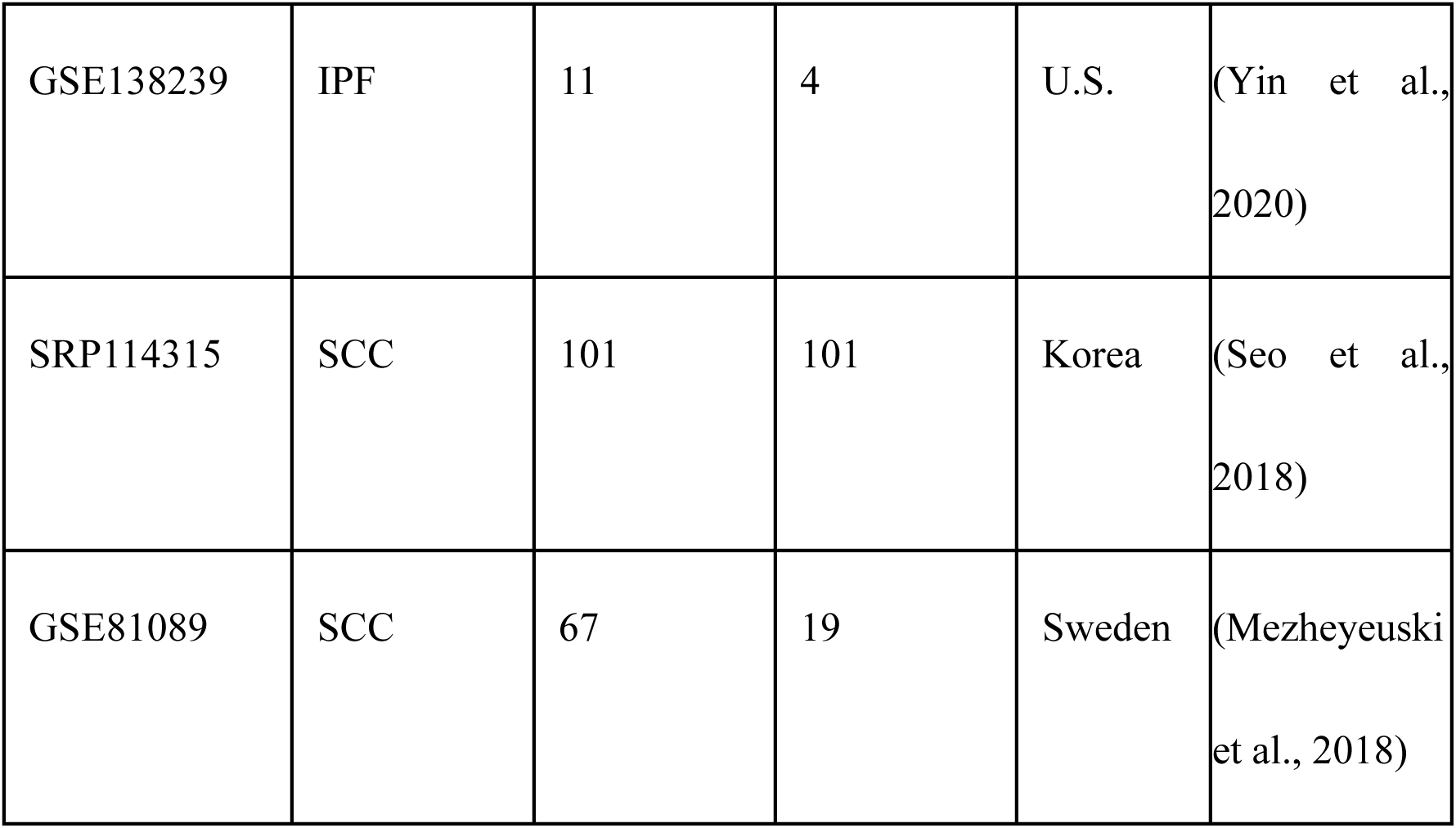
Public RNA sequencing data used in this study.

Subsequently, the transformed values were fed into the ComBat_seq function of the sva package (version 3.46.0) in R to perform BEC among the datasets. The outcomes of BEC were estimated using UMAP via the umap function of the umap package (version 0.2.10.0). UMAP visualization was conducted using the ggplot function of the ggplot2 package (version 3.4.4). After BEC, normal samples were excluded for subsequent analyses.

### Construction of ML models and feature importance

We selected the dataset with the largest sample size for each disease and designated it as the training data, whereas the remaining datasets were assigned to test data. The oversampling and downsampling of the training data were conducted using the synthetic minority oversampling technique (SMOTE) function of the DMwR package (version 0.4.1) in R. Classification models were constructed using the train function of the caret package (version 6.0-94). The training samples were split into training and validation data at a ratio of 8:2 and trained using 5-fold cross-validation. The following algorithms were selected to construct the classification models: *k*-nearest neighbors (knn), support vector machines (SVM) with radial basis function kernel (svmRadial), SVM with linear kernel (svmLinear), eXtreme gradient boosting (xgbTree), and random forest (rf). The hyperparameter settings for each ML algorithm in the grid search are detailed in Supplementary Table 1. A random restart was performed with seed values ranging from 1 to 2000 in increments of 1. Metrics of accuracy, area under the curve (AUC), and kappa value were calculated using the confusionMatrix function of the caret package.

### Somatic mutation analysis using RNA-Seq data

Somatic mutation analysis was conducted using SRP114315 samples, partially referencing the GATK Best Practices workflow for RNA-Seq data (https://github.com/broadgsa/gatk/blob/master/doc_archive/methods/Calling_variants_in_RNAseq.md). Read trimming was performed using the same method as described previously. Trimmed reads were mapped to the hs37d5 reference human genome from the 1000 Genome Project (https://www.internationalgenome.org/) using multi-sample 2-pass mapping of STAR version 2.7.0b (https://github.com/alexdobin/STAR?tab=readme-ov-file)(Dobin et al., 2013) with the --outFilterMismatchNmax 2 option, generating BAM files. These files were input into MarkDuplicates of Picard (version 2.21.8, https://broadinstitute.github.io/picard/) to identify and tag duplicate reads. The marked BAM files were sorted using Picard’s SortSam with the SORT_ORDER=coordinate option. Finally, these sorted BAM files were input into SplitNCigarReads of GATK version 4.1.4.1 (https://gatk.broadinstitute.org/) to split reads that contain Ns in their cigar string. The sorted BAM files were input into GATK’s BaseRecalibrator to generate a recalibration table for Base Quality Score Recalibration (BQSR). The BQSR and BAM files were then input into GATK’s ApplyBQSR to apply the BQSR results. The final BAM files were input into GATK’s Mutect2 to call somatic mutations, generating variant call format (VCF) files for each sample. SnpEff version 4.3t (https://pcingola.github.io/SnpEff/) was used to perform variant annotation and impact prediction in VCF files, and variants with high or moderate impact were filtered through SnpSift (version 4.3t). The selected VCF files were then converted into mutation annotation format (MAF) files using vcf2maf (version v1.6.17). Finally, the MAF files were integrated using the merge_mafs function of the maftools packages (version 2.10.05) in R.

### Network construction

Network construction involved two processes. The first process detected 13,538 mutated genes from 101 MAF files. After filtering out genes with a mutation frequency of <50% across all samples, 727 genes remained. Of these, 26 genes were identified as genes encoding TFs using the Human Transcription Factors, a public database (http://humantfs.ccbr.utoronto.ca/index.php) (Lambert et al., 2018).

In the second process, we examined the TFs that regulate the expression of 20 genes identified via feature importance using ChIP-Seq data of ENCODE (Consortium, 2011) downloaded from Harmonizome 3.0 (https://maayanlab.cloud/Harmonizome/). A total of 22,819 downstream genes were present in ENCODE ChIP-Seq data.

## Results

### Batch effect correction among multiple datasets

To detect differences in gene expression levels between IPF and SCC, we constructed ML models using public RNA-Seq datasets. We focused on protein-coding genes, long non-coding RNAs (lncRNA), and microRNAs (miRNA), which are known to be involved in lung cancer and IPF development (Ali et al., 2022; Hadjicharalambous and Lindsay, 2020; Schmitt and Chang, 2016; Wang et al., 2015, 2020). The ML model construction strategy is depicted in Fig. 1. BEC were necessary because of the use of multiple datasets deposited from different countries (Table 1). Using uniform manifold approximation and projection (UMAP), pre-BEC training and test data were plotted for each dataset (Supplementary Fig. 1), and post-BEC data were plotted for each sample category (Fig. 2). After BEC, normal and diseased samples were separated in both the training and test data, with a more distinct separation in the test data (Fig. 2).

**Figure 1.**
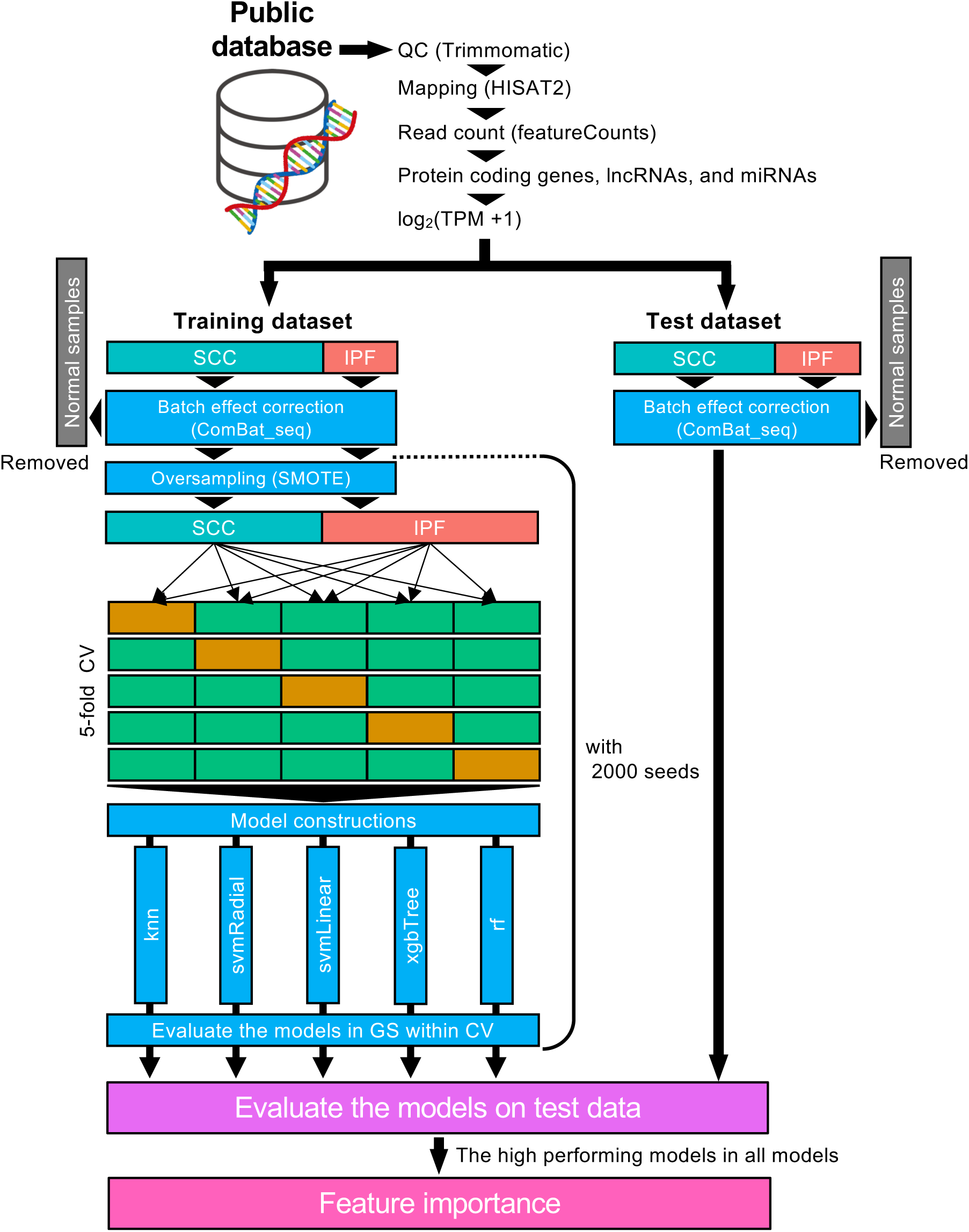
Overview of machine learning (ML) model construction and validation using test data. RNA sequencing data were obtained from public databases for both training and testing datasets. Subsequently, data were analyzed and subjected to batch effect correction using Combat_seq. The training data imbalance was addressed using the synthetic minority oversampling technique (SMOTE), and models were constructed using five algorithms through 5-fold cross-validation and grid search. From SMOTE to grid search cross-validation, 2000 seed values were utilized. Model accuracies were evaluated using the test data, and the feature importance of the highest-performing model was calculated. QC: quality control; GS: grid search; CV: cross-validation; SCC: squamous cell carcinoma; IPF: idiopathic pulmonary fibrosis.

**Figure 2.**
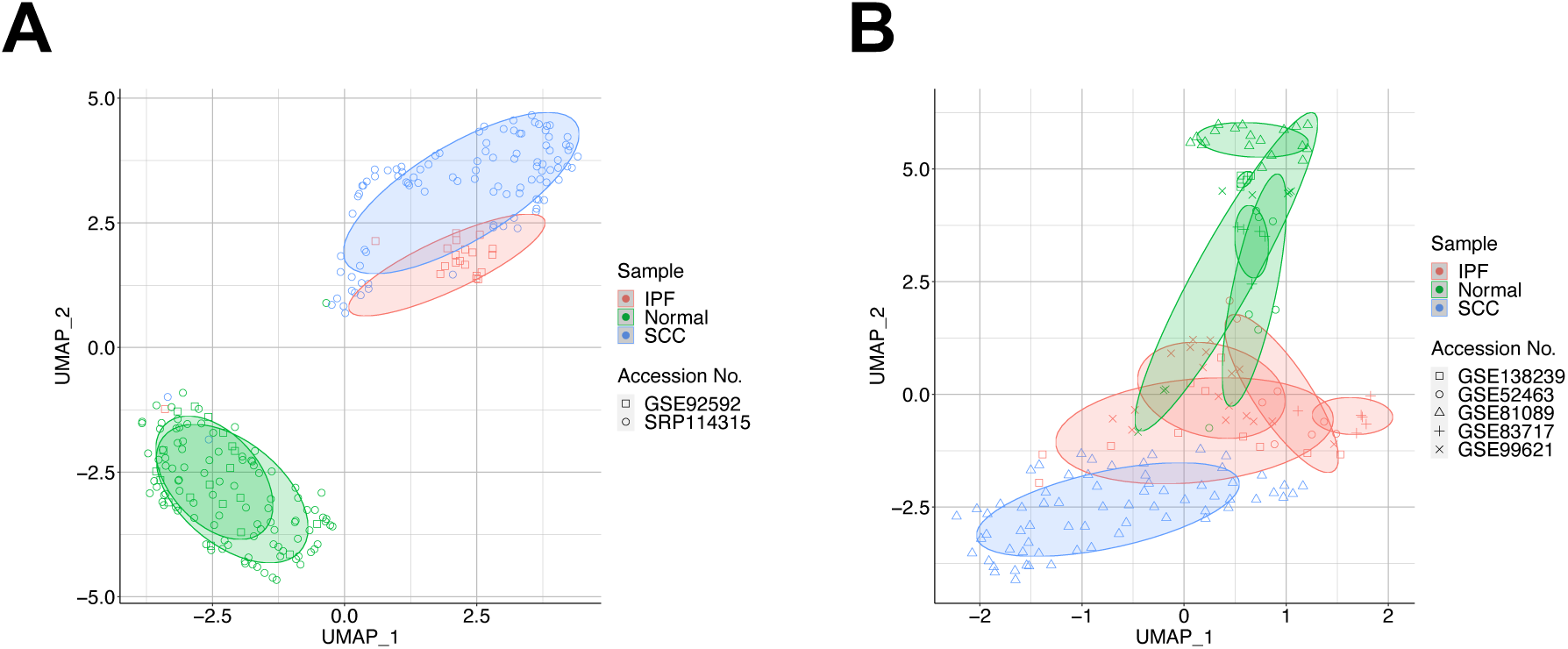
Uniform manifold approximation and projection (UMAP) of training and test data after batch effect correction (BEC). Plot colors indicate sample categories, whereas plot shapes indicate accession numbers of the public RNA sequencing data. IPF: idiopathic pulmonary fibrosis; SCC: squamous cell carcinoma. **(A)**; Training data, **(B)**; Test data.

### Construction of ML models

Biomedical datasets for rare diseases such as IPF are often severely imbalanced, rendering most ML algorithms unsuitable (Mirza et al., 2019). To address this, the SMOTE can create synthetic minority class samples (Chawla et al., 2002). We used SMOTE for oversampling IPF data and downsampling SCC data due to the extreme class imbalance in the training data. We then constructed classification models using five algorithms to classify IPF or SCC with 2000 different seed values based on the known impact of seed values on the performance of ML models (Goldberg, 2017). Optimal hyperparameters were searched during the learning process using grid search and were evaluated via 5-fold cross-validation (Supplementary Fig. 2). The model with the highest performance was selected for each seed value. Final performance evaluations were conducted based on kappa values, given the test data imbalance, generating various kappa values from five classification models (Fig. 3A). The distribution patterns of the kappa values of svmLinear and xgbTree were similar, while those of others were not (Fig. 3B). The mean and median kappa values of svmLinear and xgbTree were high among the five algorithms (Table 2), with knn showing the third highest performance. The performances of svmRadial and rf were unsatisfactory (Fig. 3A, B, and Table 2). The highest kappa values of knn, svmRadial, svmLinear, xgbTree, and rf were 0.7670, 0.4909, 0.7630, 0.7855, and 0.6383, respectively (Table 2). The accuracy, AUC, and receiver operating characteristic (ROC) curve of each model with the highest kappa value are shown in Table 3 and Fig. 3C. The accuracy, kappa, and AUC in 5-fold cross-validation of each model with the highest kappa value were ∼1 in each fold and each algorithm (Supplementary Fig. 3). According to Landis *et al*., a kappa value of 0.61–0.80 is considered “substantial” and 0.81–1.00 is considered “almost perfect” (Landis and Koch, 1977). Thus, we selected knn, svmLinear, and xgbTree models with the highest kappa values close to 0.8 for downstream analysis.

**Figure 3.**
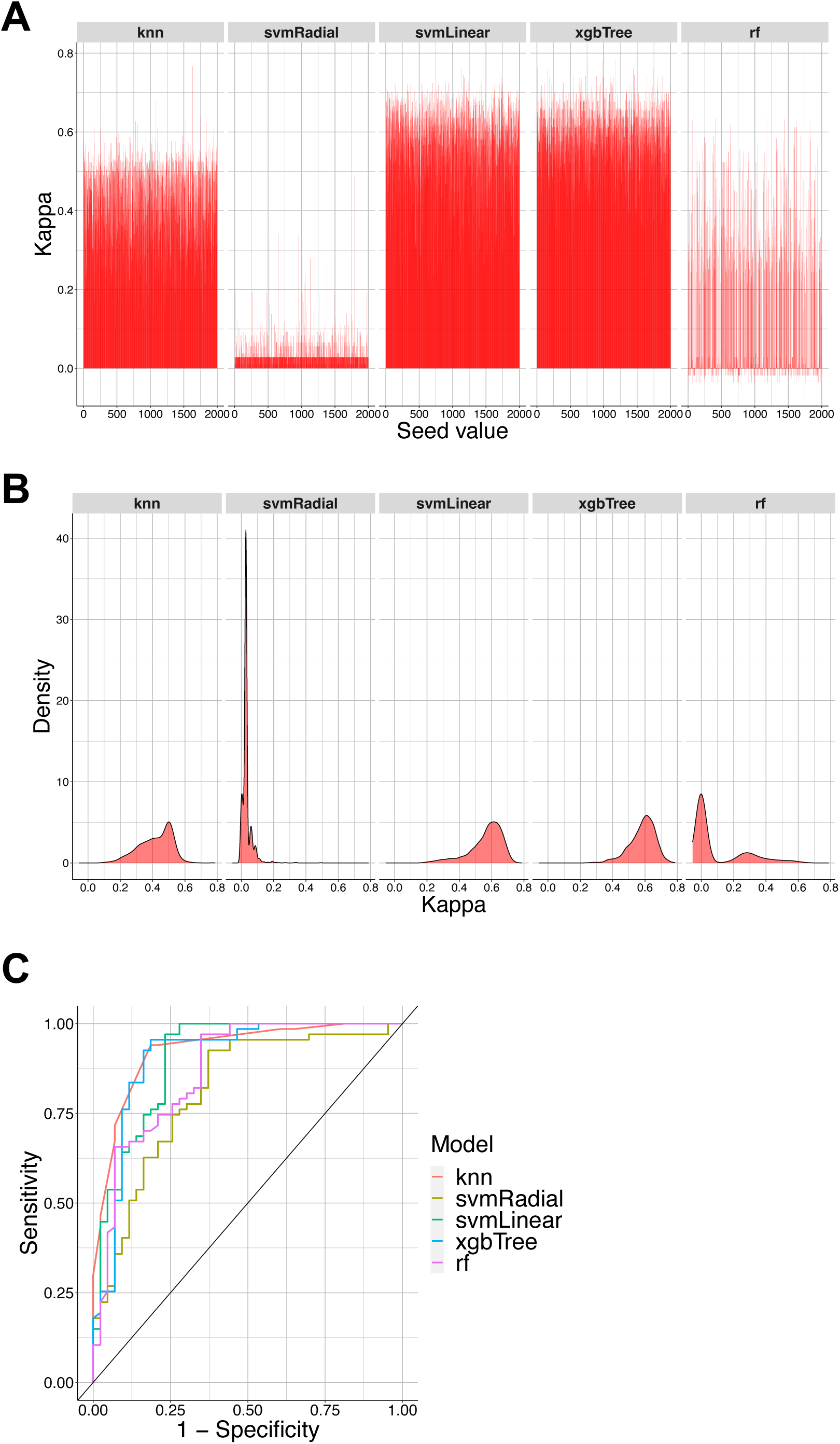
Model performance in each algorithm validated using test data. **(A)** Model kappa values in each algorithm according to seed values. **(B)** Density of kappa values in each algorithm. **(C)** ROC curve of the best-performing model in each algorithm.

**Table 2.**
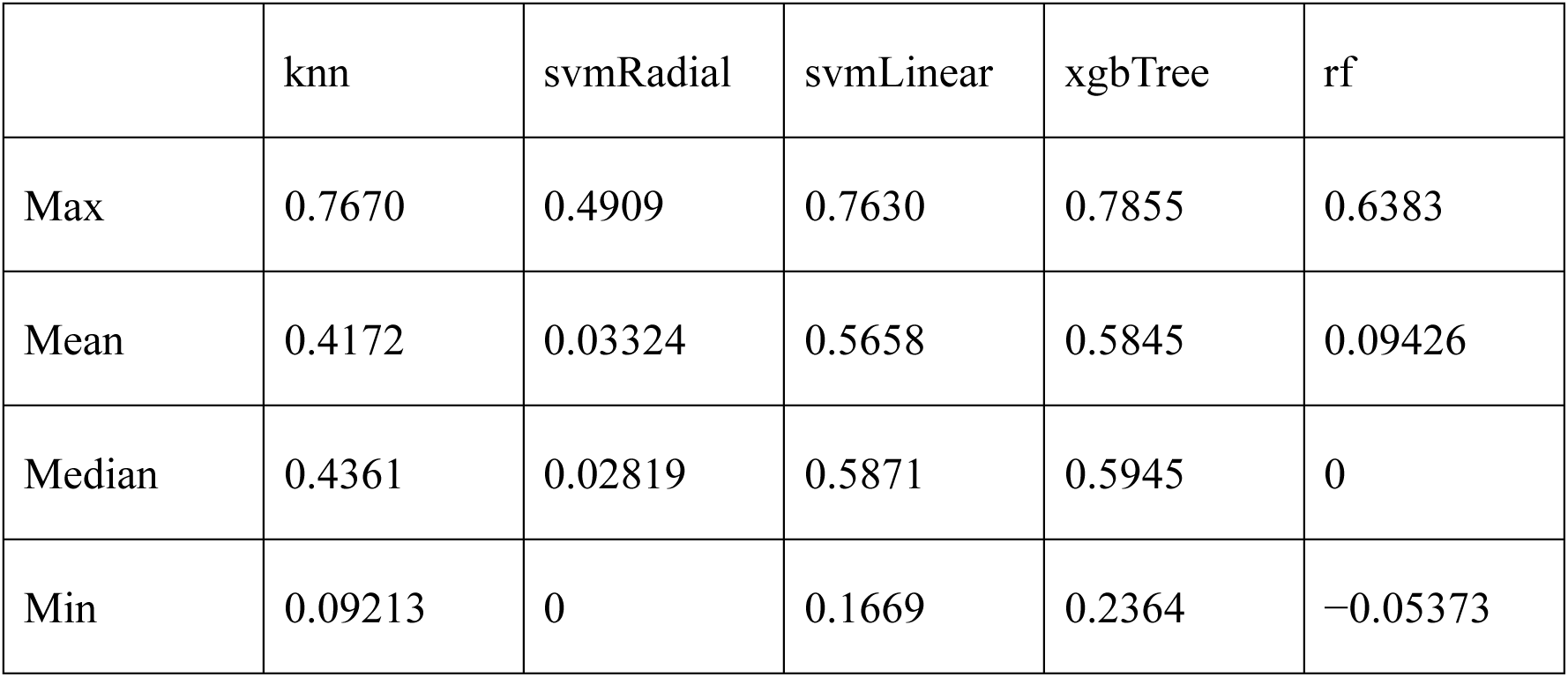
Kappa value statistics in each algorithm validated against test data.

|  | knn | svmRadial | svmLinear | xgbTree | rf |
| --- | --- | --- | --- | --- | --- |
| Max | 0.7670 | 0.4909 | 0.7630 | 0.7855 | 0.6383 |
| Mean | 0.4172 | 0.03324 | 0.5658 | 0.5845 | 0.09426 |
| Median | 0.4361 | 0.02819 | 0.5871 | 0.5945 | 0 |
| Min | 0.09213 | 0 | 0.1669 | 0.2364 | -0.05373 |

**Table 3.** Accuracy, area under the curve (AUC), and kappa value of the best-performing model in each algorithm validated against test data.

|  | knn | svmRadial | svmLinear | xgbTree | rf |
| --- | --- | --- | --- | --- | --- |
| Accuracy | 0.8909 | 0.7636 | 0.8909 | 0.9000 | 0.8364 |
| AUC | 0.9271 | 0.8119 | 0.9045 | 0.9065 | 0.8698 |
| Kappa | 0.7670 | 0.4909 | 0.7630 | 0.7855 | 0.6383 |

### Feature importance

Significant genes were identified from the classification models (knn, svmLinear, and xgbTree) based on feature importance. The top 10 genes included *MAGEA4* in all models; *SCARA5*, *PLA2G2A*, *ENSG00000234638*, *HPSE2*, *ENSG00000279712*, *DES*, and *PRUNE2* in knn and svmLinear; and *PRG4* in svmLinear and xgbTree. *ENSG00000250920* and *PZP* were exclusive to knn; *LINC00602* to svmLinear; and *LINC02990*, *MAFA-AS1*, *VCX2*, *S100A7A*, *GPR50*, *ENSG00000272715*, *HOXD11*, and *TMPRSS11B* to xgbTree (Fig. 4A). The scaled importance of the top 10 genes was >90 in knn and svmLinear (Fig. 4B, C) and ranged from 2.83 to 100 in xgbTree (Fig. 4D). Compared with the normal tissue, the expression levels of the abovementioned 20 genes were drastically altered in SCC lung tissue, with slight alterations in IPF compared with SCC (Fig. 4E).

**Figure 4.**
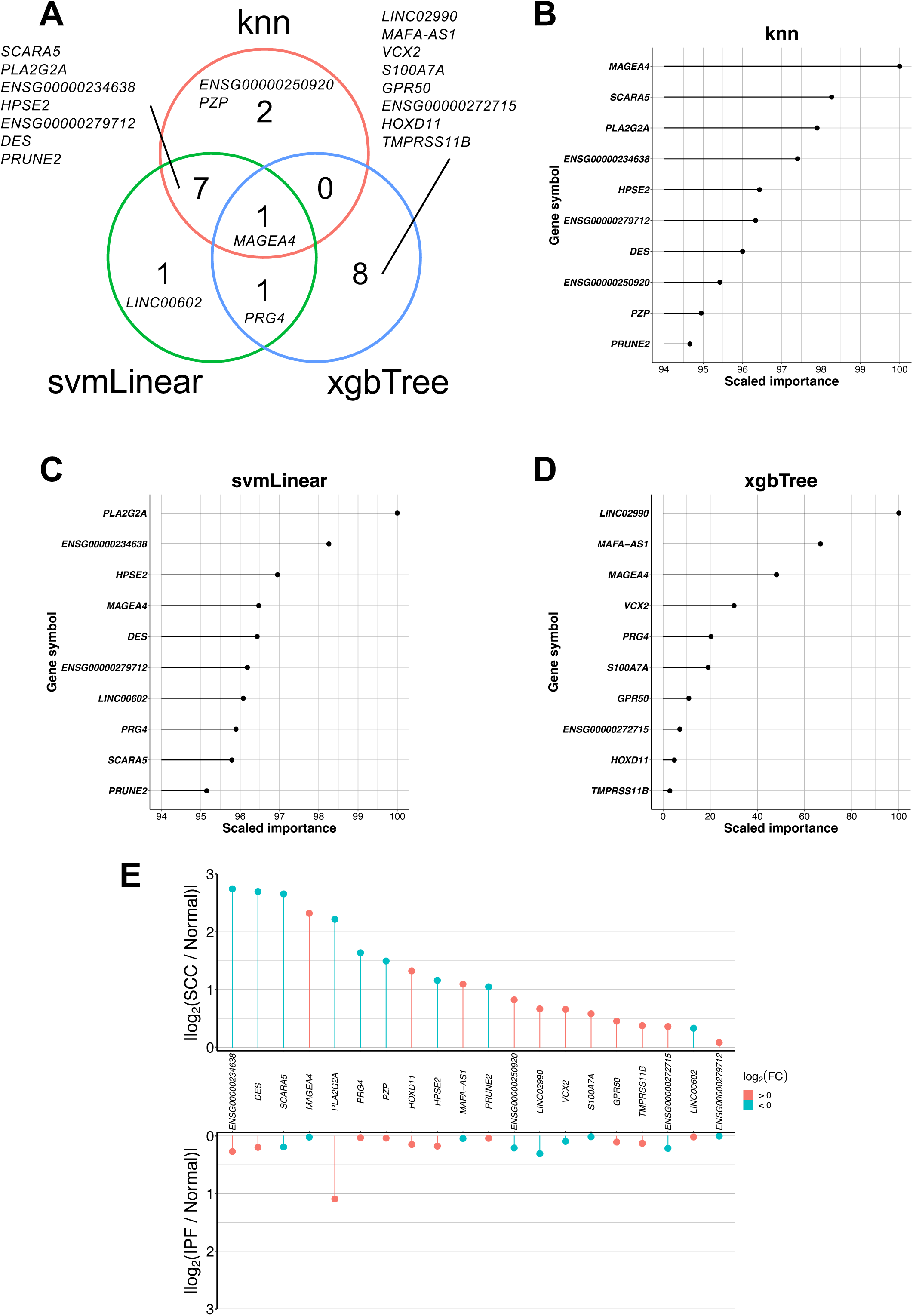
Identification of significant genes based on feature importance. **(A)** Overlap among the top 10 genes based on feature importance in knn, svmLinear, and xgbTree. Top 10 genes based on feature importance in **(B)** knn, **(C)** svmLinear, and **(D)** xgbTree. **(E)** Log_2_(FC) values of the top 20 differentially expressed genes between normal and IPF, or normal and SCC. Red and blue indicate positive and negative values, respectively. FC; fold change

### Somatic mutation analysis of transcription factors regulating the expression of genes identified via feature importance

Intratumor heterogeneity refers to cellular variations within tumors (Ono et al., 2021), with unique genomic profiles observed even in adjacent tumors of the same patient; this phenomenon is observed in SCC (de Bruin et al., 2014). This makes it challenging to correlate gene expression from one tumor sample with somatic mutation data from an adjacent sample. Therefore, although the dataset with accession no. SRP114315 includes both RNA-Seq and whole-exome sequencing (WES) data, we used the RNA-Seq data to identify genomic profiles causing gene expression differences, as the tumor tissue samples for RNA-Seq and WES were different (Seo et al., 2018). Past genome-wide sequencing studies have detected a higher number of somatic mutations in non-small cell lung cancer (NSCLS) than in other cancers, indicating a significant role of these mutations in NSCLS onset (Vogelstein et al., 2013). Therefore, we investigated somatic mutations affecting gene expression, focusing on mutations in genes encoding transcription factors (TFs). The identification process, as shown in Fig. 5, involves two processes. After the first process, variant classification revealed that frame shift insertions were the most common mutations, followed by missense mutations with a median of 32 variants per sample (Fig. 6A, B). Twenty-six TFs with mutations in >50% of samples are shown in Fig. 6C and Supplementary Fig. 4. After the second process, 13 downstream genes were found to overlap with the important feature genes detected by the ML models, and according to chromatin immunoprecipitation sequencing (ChIP-Seq) data from the encyclopedia of DNA elements (ENCODE) data, these genes were regulated by one or more of the 128 TFs (Fig. 7A, Supplementary Data 1).

**Figure 5.**
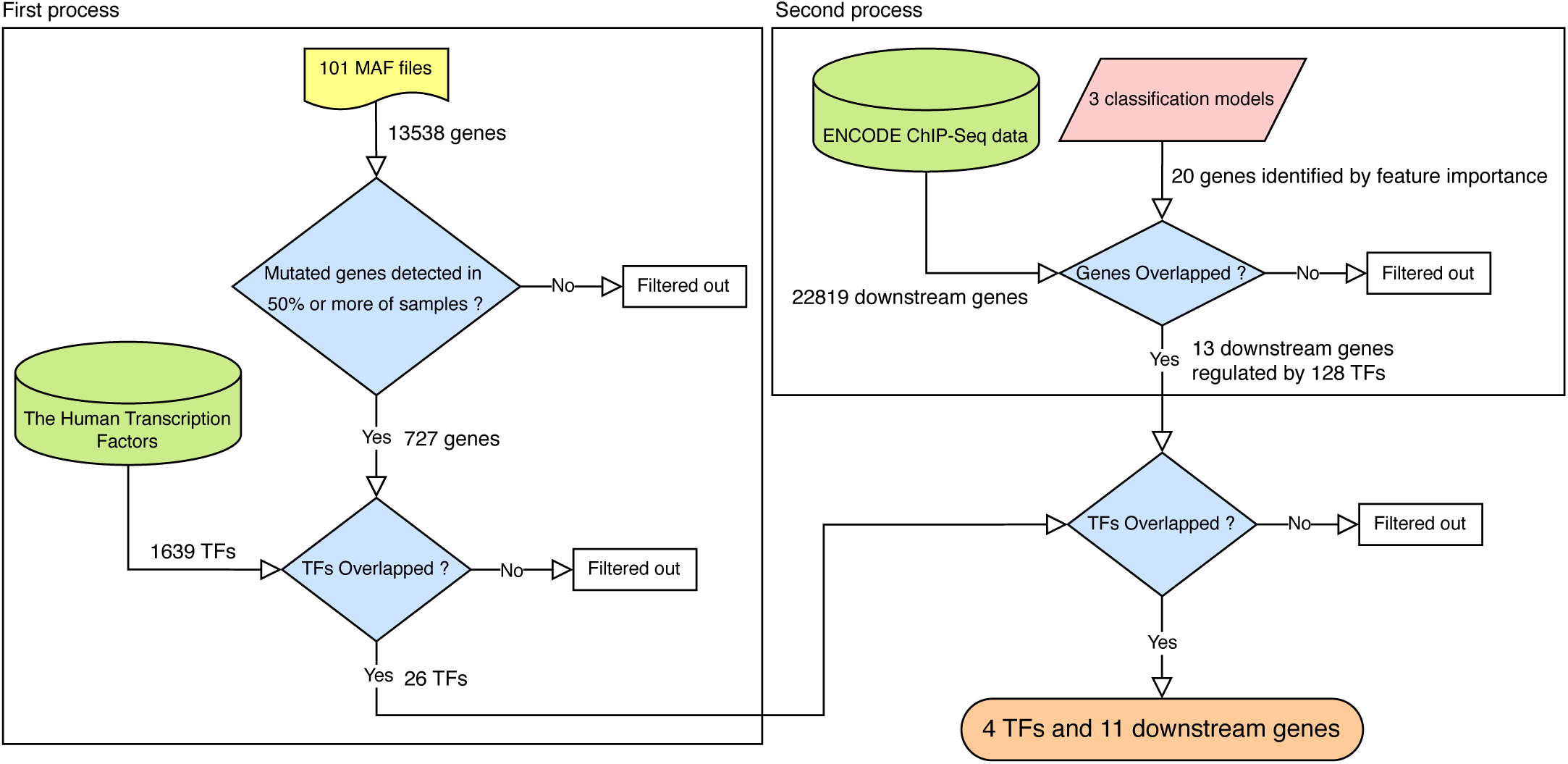
Flowchart for the identification of somatic mutations in transcription factors (TFs) that regulate the expression of genes identified via feature importance. Diagram was generated using draw.io (https://app.diagrams.net/).

**Figure 6.**
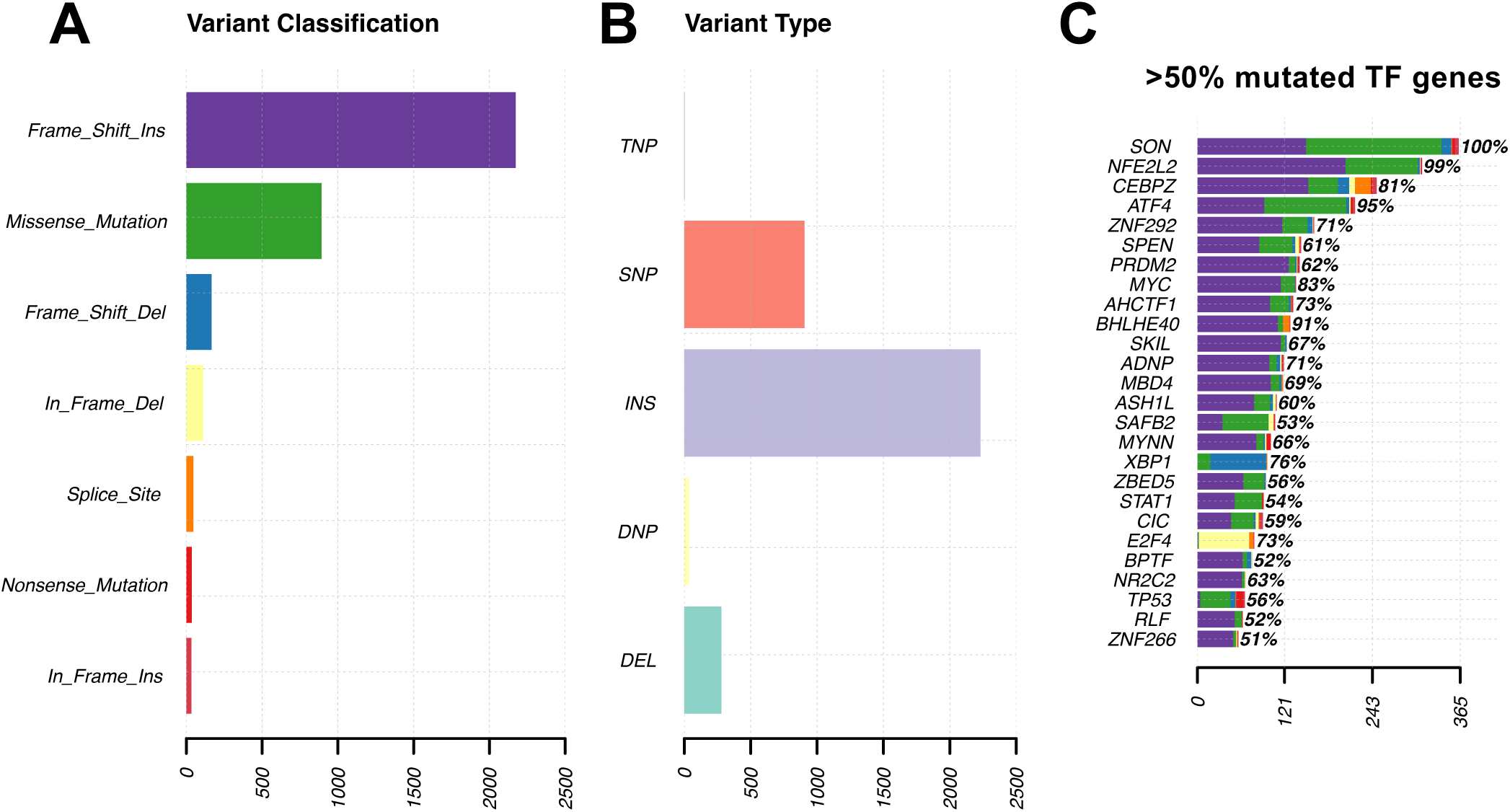
Summary of somatic mutation analysis in squamous cell carcinoma (SCC) samples. **(A)** Variant classification and **(B)** variant type. In both cases, the horizontal axis represents the number of variants detected across all samples. **(C)** List of TFs for which a mutation was detected in >50% of samples. Horizontal axis represents the number of variants detected for each TF. Plots were constructed using the plotmafSummary function in the maftools package (version 2.10.05) in R.

**Figure 7.**
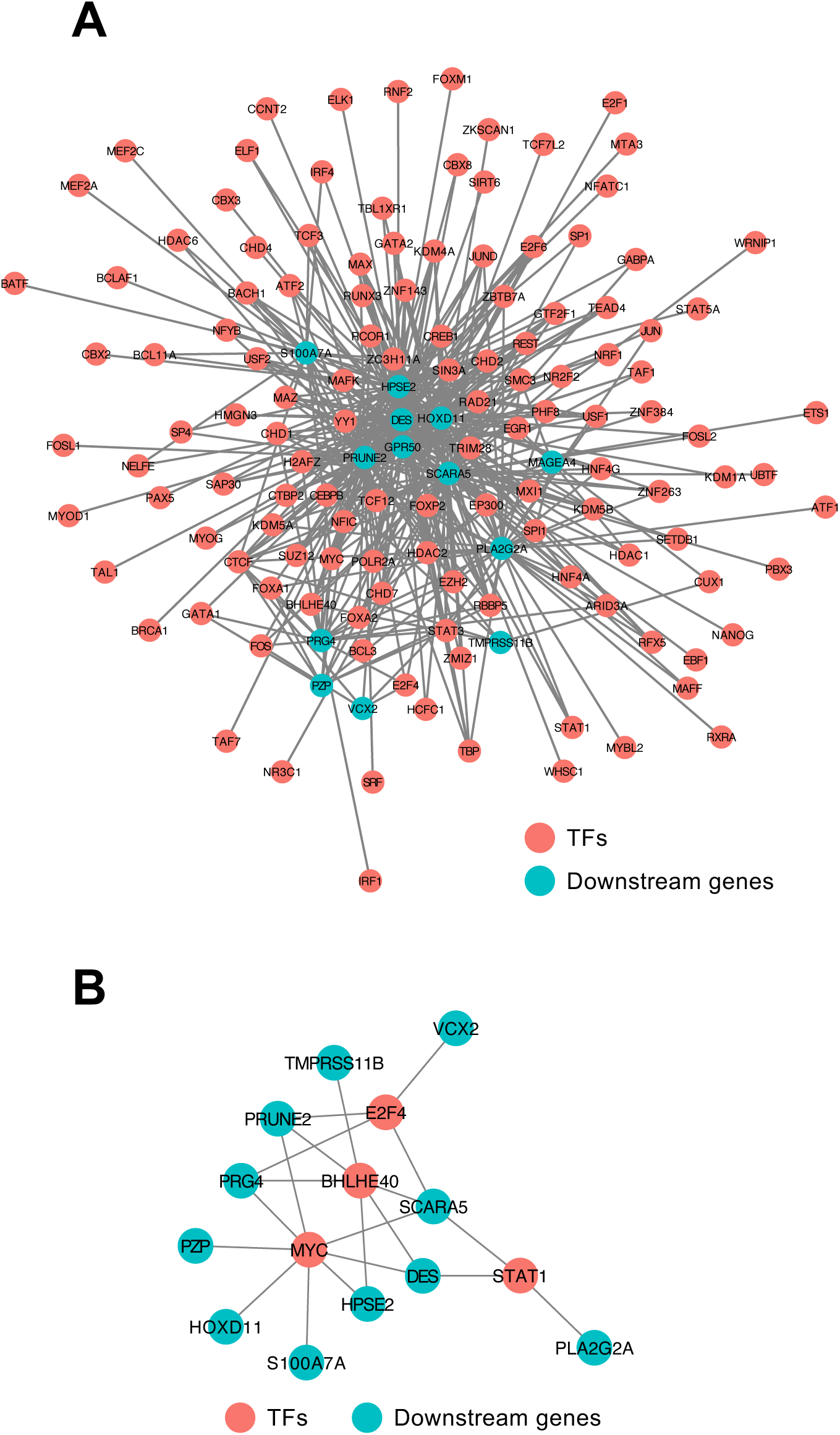
Molecular networks consisting of transcription factors (TFs) with somatic mutations and the genes regulated by these TFs. **(A)** A network of 13 downstream genes identified based on the second process and 128 TFs that regulate these genes. **(B)** A network of four TFs identified based on the first and second processes along with the genes regulated by these TFs according to feature importance. Red and blue nodes represent TFs and downstream genes, respectively. Connections (edges) indicate regulatory relationships between TFs and downstream genes. Networks were visualized using Cytoscape version 3.10.0 (https://cytoscape.org/).

The TFs *MYC*, *BHLHE40*, *STAT1*, and *E2F4* were identified as overlaps between the 26 TFs from the first process and 128 TFs from the second process. All detected somatic mutations in these four TF genes are documented in Supplementary Data 2. In *MYC*, *BHLHE40*, and *STAT1*, Frame_Shift_Ins mutations were most frequently detected, followed by Missense_Mutation (Fig. 6C and Supplementary Fig. 4). In *E2F4*, the majority of somatic mutations were In_Frame_Del (Fig. 6C and Supplementary Fig. 4). The most common mutations in *BHLHE40* and *E2F4* were not located in the domain region (Supplementary Fig. 5A, C). In contrast, the most frequent mutations in *MYC* and *STAT1* were located in the HLH domain and STAT_alpha domain, respectively (Supplementary Fig. 5B, D). Less frequent mutations were scattered throughout the entire sequence (Supplementary Fig. 5A–D). Based on ENCODE data, the expression of 11 downstream genes was regulated by one or more of these four TFs (Fig. 7B). In addition, missense mutations of these four TFs were detected in at least one of the 14 samples in the TCGA-LUSC dataset (Supplementary Fig. 6).

## Discussion

In this study, we constructed ML models to categorize patients into IPF or SCC using RNA-Seq data from seven different datasets. The advantage of merging gene expression data batches to enhance statistical power is often compromised by batch effects. These effects are unwanted data variations caused by technical differences across batches (Zhang et al., 2020). Therefore, it is crucial to minimize these batch effects, particularly in the training data. The training data after BEC were distinctly distributed for each sample category, indicating successful correction of batch effects. Although the test data were more widely spread than the training data, normal and diseased samples were separately distributed, indicating successful correction of batch effects.

ML methods are particularly useful when dealing with data where the number of samples (*n*) is considerably greater than the number of features (*p*). However, in omics data, the scenario is reversed, with the number of features (such as genes and proteins) significantly exceeding the number of samples, leading to a data structure expressed as *n* << *p* (Teschendorff, 2019). This can result in overfitting when applying ML methods to omics data. Overfitting occurs when the derived predictive model fits a random variation in the data that does not represent the true biological variation associated with the phenotype of interest (Simon et al., 2003; Teschendorff, 2019). Therefore, it is crucial to validate the performance of ML models using test data that are independently constructed from the training data when conducting ML with omics data (Teschendorff, 2019). In the process of training complex networks, employing various random seeds can result in different final solutions, each with its own accuracy (Goldberg, 2017). This method is known as random restarts (Goldberg, 2017). In a random restart, the algorithm begins with several initial solutions (random initial states) and conducts the optimization process for each solution. In other words, the same problem is tackled multiple times under various initial conditions, and the optimal solution is selected based on the outcomes of each attempt. This method’s advantage lies in its ability to mitigate the risk of local optima entrapment, enabling a more extensive solution search by testing multiple initial solutions. However, this could increase computational costs, necessitating an efficient implementation. Therefore, it is advisable to execute the training process multiple times with varying random seeds when computational resources allow, followed by the selection of the random seed that performs best on the development sets (Goldberg, 2017). In this study, we employed the random restart technique to select the high-performing classification model. The classification models exhibited high performance in 5-fold cross-validation, indicating potential overfitting. Therefore, independently constructed test data were used for evaluation with 2000 seed values. The svmLinear, knn, and xgbTree classification models were selected based on their kappa values when validated against this test data. The kappa values for all three models exceeded 0.75, which, according to the report by Landis *et al*. (Landis and Koch, 1977), indicates substantial model performance. This demonstrates the feasibility of constructing high-performing ML models, even for omics data with *n* << *p*, when independently constructed test data and random restarts technique are available. As mentioned above, the classification model constructed in this study consistently exhibited high performance during grid search, regardless of hyperparameter conditions, indicating that random restarts are more critical and warrant greater consideration than model-specific hyperparameters in studies using ML for omics data with *n* << *p*. When the goal is to detect differences between two classes by extracting feature importance from the model, as in this study, selecting the seed value that yields the highest accuracy among those obtained through random restarts has proven effective in achieving a highly accurate model. In contrast, varying accuracy results may be achieved depending on the test data used, even with the same seed value. Therefore, when the goal is to construct a general-purpose ML model, the random restart technique may be inappropriate.

Interestingly, the top 10 feature importance scores calculated based on svmLinear and knn classification models all fell within the range of 90–100. In contrast, those calculated based on the xgbTree classification model showed a wide range of values, from 2 to 100, indicating a high degree of variability. This suggests that the svmLinear and knn models make predictions using a larger set of genes, while the xgbTree model relies on a smaller, more selective set of genes for its predictions. The difference likely stems from algorithmic variations. The top 10 genes with high feature importance in knn and svmLinear largely overlapped, unlike those in xgbTree. Despite this, the kappa values and accuracy of these three models were similar, suggesting that predictions can be made using two distinct gene lists.

Feature importance based on three classification models revealed 20 key genes for IPF and SCC classifications. These genes could explain the differences in gene expression levels between IPF and SCC. Considering the significant changes between SCC and normal lungs and the minor differences between IPF and normal lungs, most of these 20 genes are likely associated with carcinogenesis rather than fibrosis. Indeed, among the 20 genes, changes in the expression profiles of some of those genes have been observed in patients with SCC. According to a previous study, compared with normal samples, the promoter region of *SCARA5* showed increased methylation levels in the lung tissue of patients with SCC, leading to reduced gene expression (Peng et al., 2021). In another study, the expression of *PLA2G2A* was decreased due to somatic mutations in tumor suppressor genes (TSGs; *TP53*, *CDKN2A*, *PTEN*, *RB1*, and *BRCA1*) in the lung tissues of patients with SCC (Kim et al., 2021). In the current study, of the 101 patients with SCC, 78 had somatic mutations in at least one of the abovementioned TSGs (Supplementary Data 3). The expression of the Desmin protein, encoded by *DES*, was lower in the lung tissues of patients with SCC than in those of patients with IPF (Fallahian et al., 2018). The expression levels of *PRUNE2*, *PZP*, *PRG4*, *HOXD11*, and *MAFA-AS1* were decreased in the lung tissues of patients with SCC compared with those in normal lung tissue (Chen et al., 2023; Tian et al., 2023; Wu et al., 2020; Zhang et al., 2017). *TMPRSS11B* and *S100A7A* expression levels were elevated in SCC lung tissue compared to normal lung tissue (Chen et al., 2013; Updegraff et al., 2018). *MAGEA4* mRNA is uniquely expressed in SCC lung tissue but not in ADC and normal lung tissues (Peikert et al., 2006). VCX2, identified as a cancer/testis antigen, is also expressed in SCC lung tissue (Taguchi et al., 2014). In this study, *PLA2G2A* emerged as the only gene showing marked differences in expression levels between IPF-affected and normal lungs, with higher expression observed in patients with IPF (Bauer et al., 2015; Jaiswal et al., 2023), suggesting a link between increased *PLA2G2A* expression and fibrosis. These findings emphasize the consistency between the high feature importance genes identified in our classification models and the known pathophysiological and molecular characteristics of IPF and SCC. Thus, beyond computational accuracy measured using metrics such as kappa values, the classification model accurately distinguishes SCC from IPF biologically. Mouse pancreatic carcinoma cells implanted in *HPSE2* knockout mice show enhanced development compared with that in wild-type mice. Additionally, *GPR50* knockdown results in decreased cell proliferation and migration in hepatocellular carcinoma cell lines. These findings suggest that *HPSE2* and *GPR50* play roles in tumorigenesis suppression (Kayal et al., 2023; Saha et al., 2020). Although *HPSE2* and *GPR50* are implicated in other cancer pathologies, their association with SCC or IPF remain unelucidated. *LINC00602*, *LINC02990*, *ENSG00000272715*, *ENSG00000234638*, *ENSG00000279712*, and *ENSG00000250920* are listed as lncRNAs in the Ensembl genome database (https://www.ensembl.org/). However, no studies have reported on these lncRNAs, despite their potential role in carcinogenesis. Therefore, our study is the first to report an association between these lncRNAs and SCC. The top 20 genes, identified based on their high feature importance, were linked to carcinogenesis rather than fibrosis. Through analysis of SCC samples, somatic mutations regulating these genes were detected, with a focus on TFs. ChIP-Seq data from ENCODE revealed a connection between these 20 genes and four TFs, including *BHLHE40*, *MYC*, *STAT1*, and *E2F4*. Somatic mutations of these four TFs were identified using the TCGA-LUSC dataset, despite low mutation frequency, suggesting that they are not dataset-specific mutations. The somatic mutations in the four TF genes may influence the expression of downstream genes associated with SCC pathology, including *VCX2*, *TMPRSS11B*, *PRUNE2*, *PRG4*, *PZP*, *SCARA5*, *DES*, *HPSE2*, *HOXD11*, *S100A7A*, and *PLA2G2A11*. Therefore, such mutations, if present in the lungs of patients with IPF, could be considered a potential carcinogenic risk.

## Conclusions

In this study, we developed ML models to distinguish SCC from IPF, achieving high accuracy even with omics data where *n* << *p*. The model identified 20 differentially expressed genes between IPF and SCC, indicating their potential role in SCC pathogenesis. We also detected somatic mutations in four TFs that regulate 11 of the 20 identified genes. These discoveries enhance our understanding of the molecular mechanisms underlying SCC complications in patients with IPF and offer new perspectives for preventing SCC complications in these patients.

## Author contributions

Y.N. and K.M. conceived and designed the experiments. Y.N. performed the experiments and analyzed the data. Y.N. contributed to the writing of the draft manuscript. K.M. reviewed and edited the draft manuscript. All authors discussed the data and the manuscript. All authors have read and approved the final manuscript.

## Acknowledgments

We thank the members of the Artificial Intelligence Center for Health and Biomedical Research in the National Institutes of Biomedical Innovation, Health and Nutrition for their valuable support and discussion.

