## Supplementary Information for "Identification of the differences in molecular networks between idiopathic pulmonary fibrosis and lung squamous cell carcinoma using machine learning"

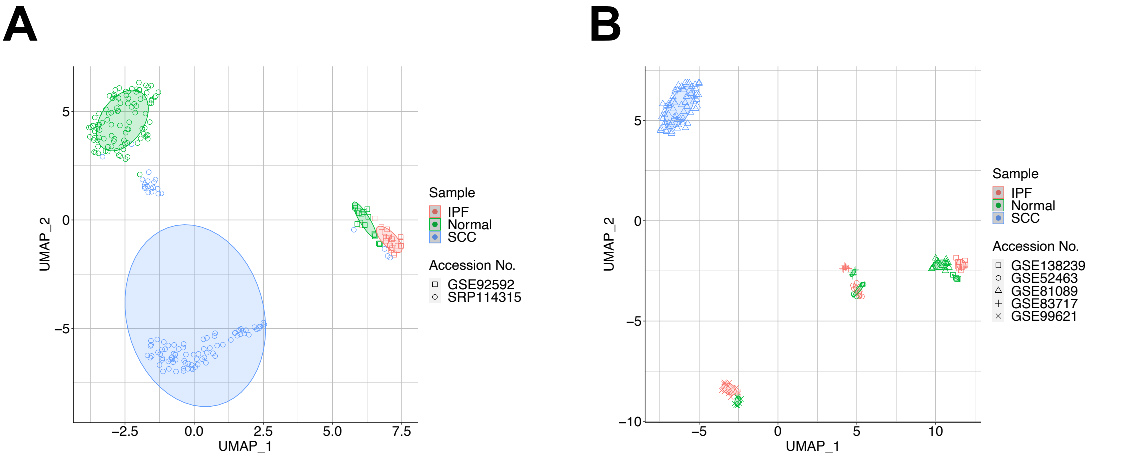


Supplementary Fig. 1 UMAP of training (**A**) and test data (**B**) before BEC. Plot colors indicate sample categories, and plot shapes represent accession numbers of the public RNA sequencing data.

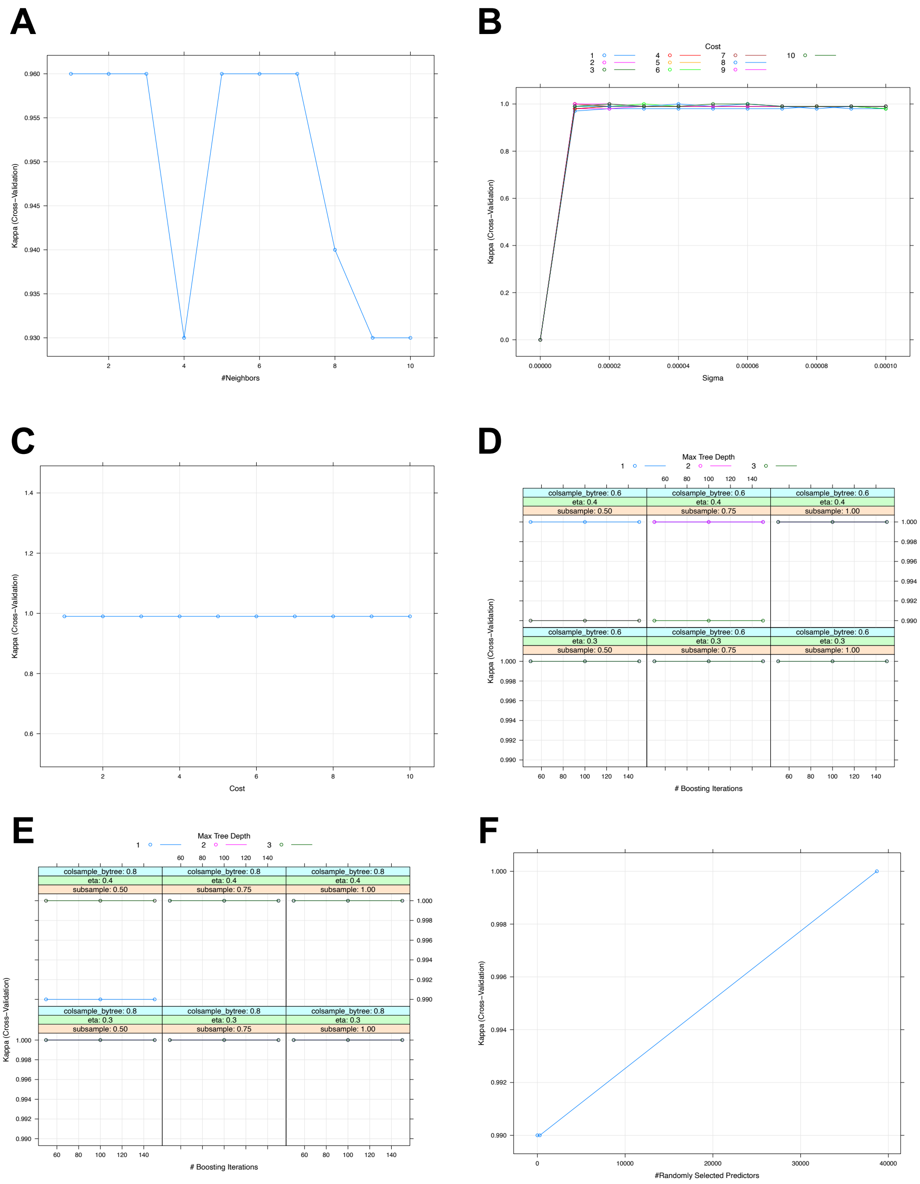


Supplementary Fig. 2 Kappa value of each hyperparameter setting during grid search cross-validation. **A**) *k*-nearest neighbors (knn), **B**) Support vector machines (SVM) with radial basis function kernel (svmRadial), **C**) SVM with linear kernel (svmLinear), **D and E**) eXtreme gradient boosting (xgbTree), **F**) random forest (rf).

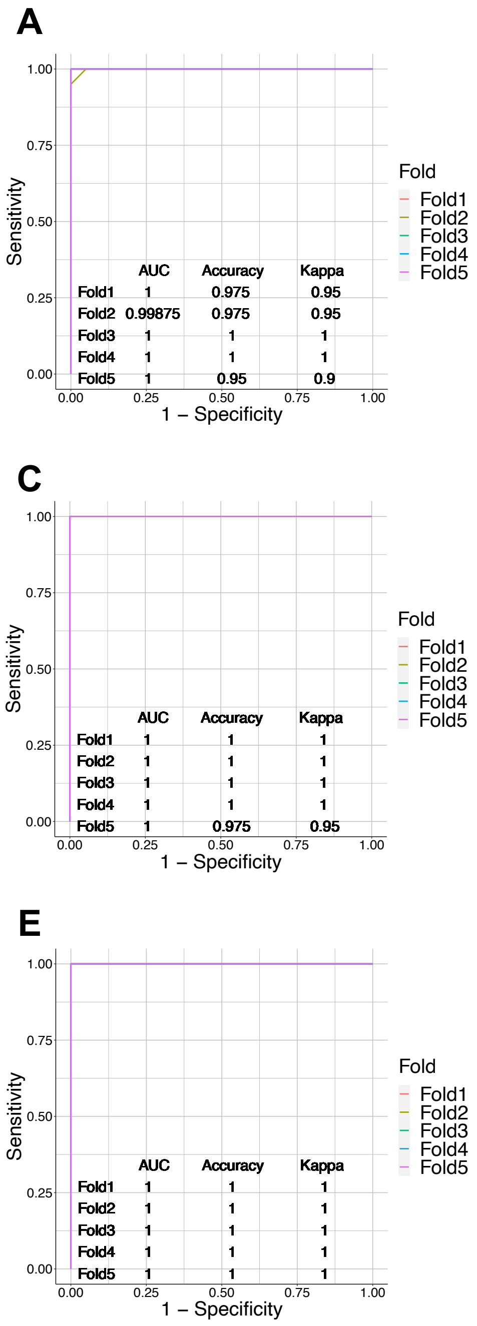


Supplementary Fig. 3 The ROC curve, AUC, accuracy, and kappa value of each fold in 5-fold cross-validation. **A**) knn, **B**) svmRadial, **C**) svmLinear, **D**) xgbTree, **E**) rf.

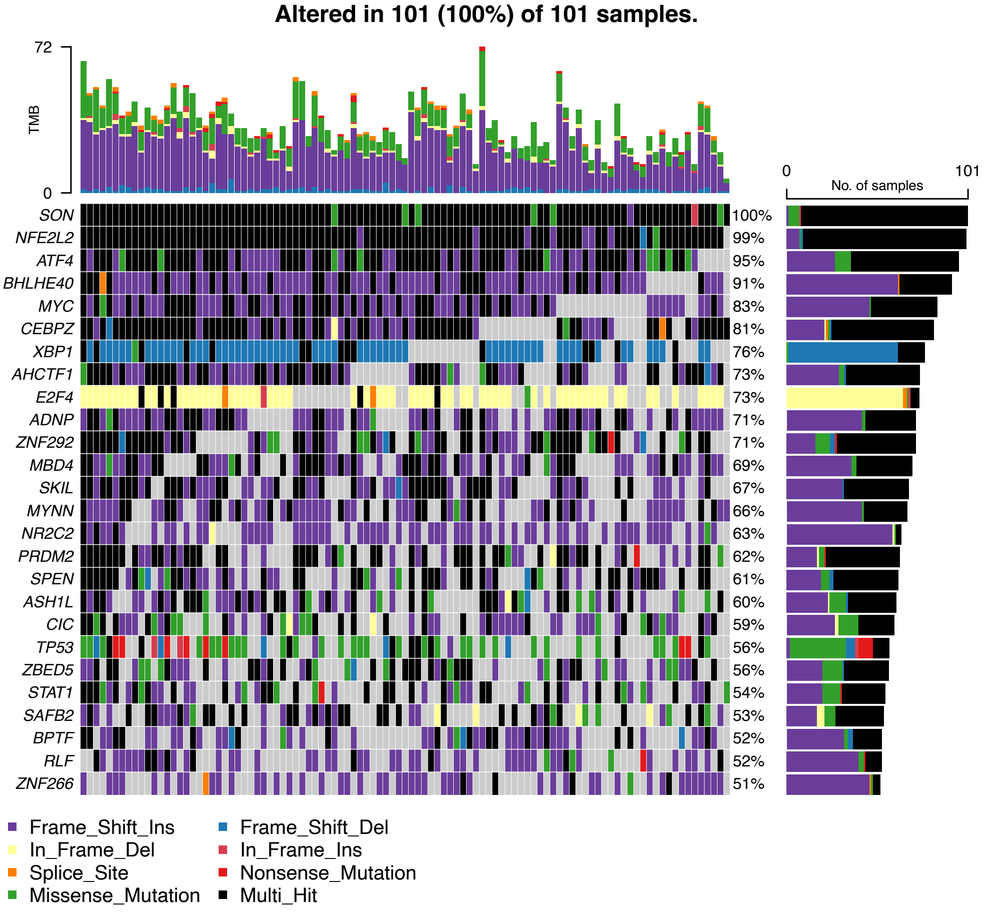


Supplementary Fig. 4 Oncoplot of somatic mutation analysis in SCC samples. The TFs with mutations in >50% of samples are shown. The plot was constructed using the oncoplot function of the maftools package (version 2.10.05) in R. TMB; tumor mutation burden.

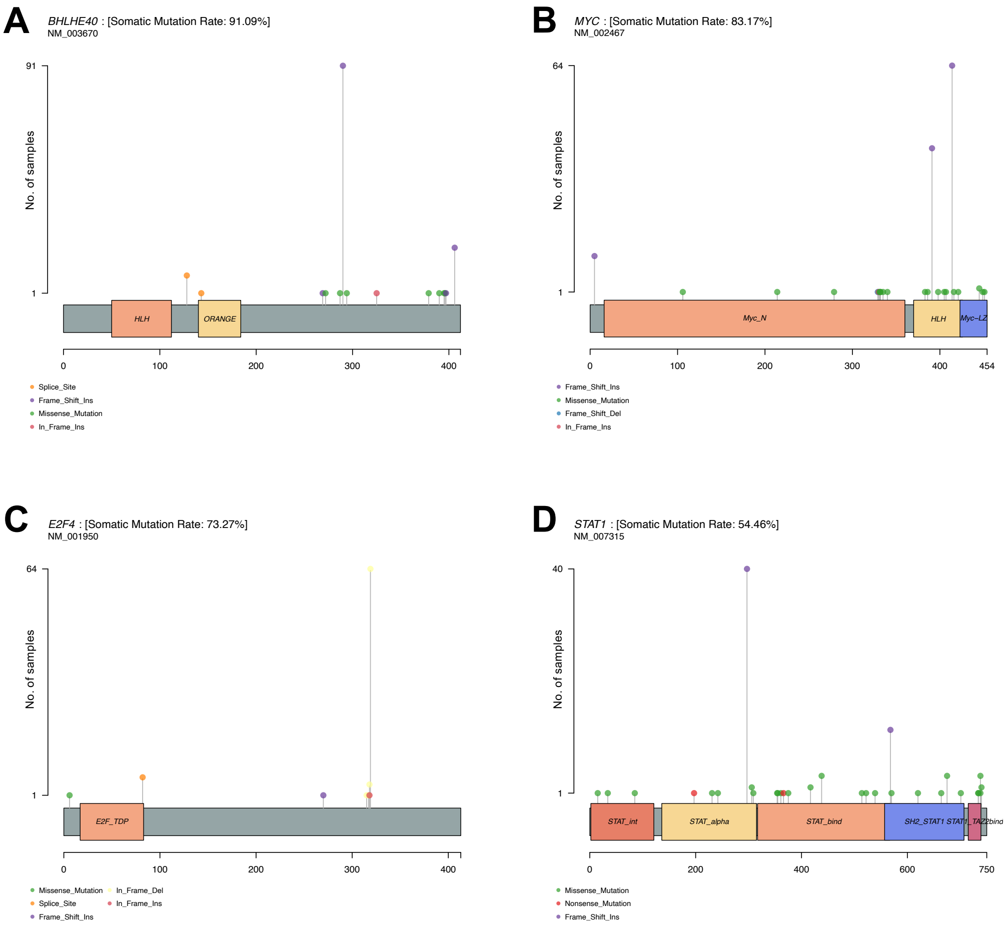


Supplementary Fig. 5 The frequency and types of somatic mutations in the genes encoding the four TFs in Fig. 7b. Lollipop plots were constructed using the lollipopPlot function of the maftools package (version 2.10.05) in R. **A**) *BHLHE40*, **B**) *MYC*, **C**) *E2F4*, **D**) *STAT1*

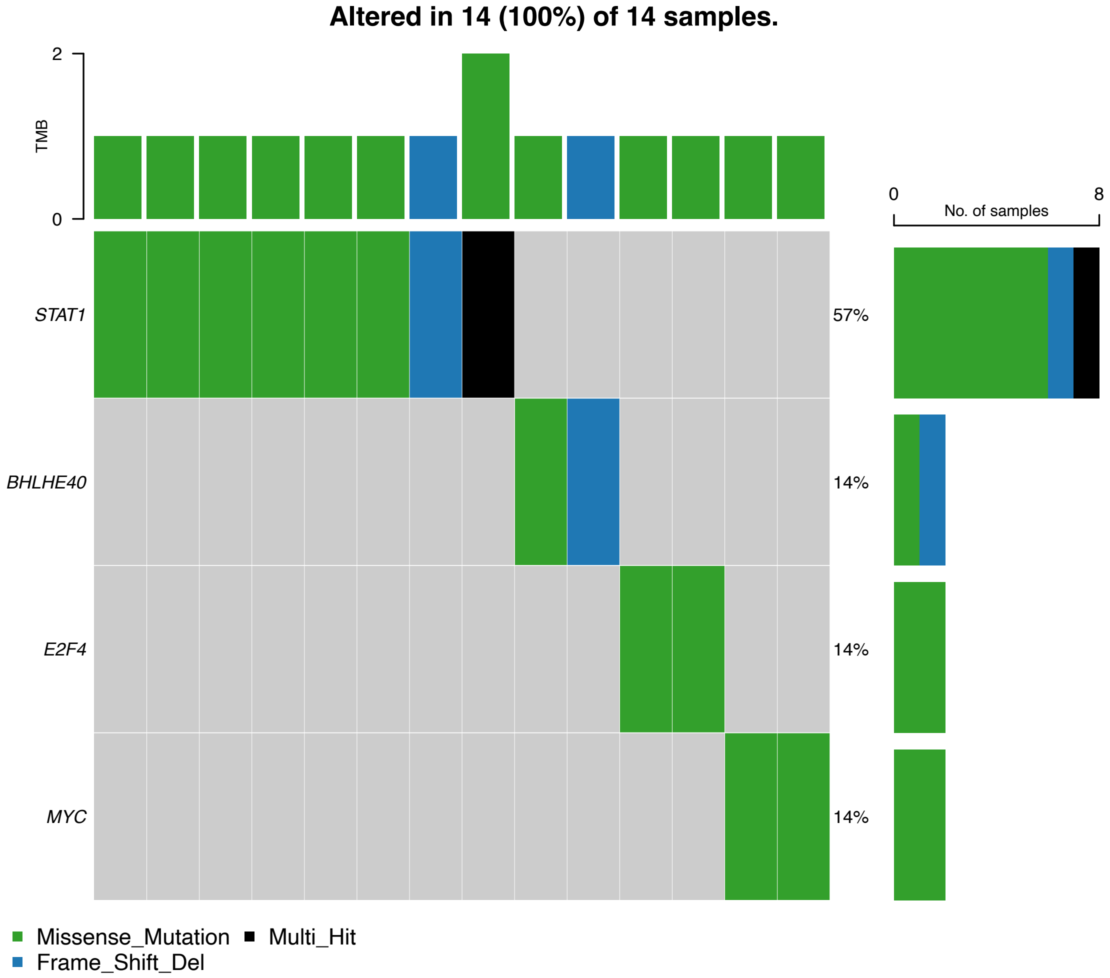


Supplementary Fig. 6 Oncoplot of somatic mutation analysis in the LUSC samples of The Cancer Genome Atlas (TCGA) dataset. The TFs *MYC*, *BHLHE40*, *STAT1*, and *E2F4* are shown. The TCGA-LUCS dataset was downloaded from the National Cancer Institute Genomic Data Commons Data Portal (https://portal.gdc.cancer.gov/v1/) using the TCGAbiolinks package (version 2.26.0) in R. The plot was constructed using the oncoplot function of the maftools package in R. TMB; tumor mutation burden.

Supplementary Table 1 Hyperparameter settings of each ML algorithm in grid search.

| knn | svmRadial | svmLinear | xgbTree | rf |
|---|---|---|---|---|
| k = 1 to 10 in 1 increment | C = 1 to 10 in 1 increment<br>sigma = 0 to 0.0001 in 0.00001 increment | C = 1 to 10 in 1 increment | Default setting | Default setting |
